# The Healthy Nevada Project: rapid recruitment for population health study

**DOI:** 10.1101/250274

**Authors:** Joseph J Grzymski, Max J Coppes, Jim Metcalf, Christos Galanopoulos, Chris Rowan, Michele Henderson, Robert Read, Harry Reed, Bruce Lipp, Dave Miceli, Susan Rybarski, Anthony Slonim

## Abstract

**Background:** Nevada ranks in the bottom half of overall health rankings in the United States. The majority of residents of Northern Nevada live in Washoe County, which is confounded with high age-adjusted death rates for heart disease, cancer and chronic lower respiratory disease.

**Methods:** Saliva as a source of DNA was collected from adults in Northern Nevada as the first phase of a much larger (100,000 participants) effort to contribute to comprehensive population health studies in Nevada. The personal genetics company 23andMe was used to genotype the first 10,250 participants and deliver their custom ancestry, traits, wellness, and carrier status reports.

**Results:** The study was announced by Governor Brian Sandoval on September 15, 2016 and within two days the registration of 9,700 volunteers for an appointment was complete. Processing of 9,344 participants was achieved in 3 months, with a no-show rate of just over 11%. The participant population was skewed to female and was less racially diverse than the population.

**Conclusion:** DNA genotyping was administered free-of-charge and the patient population was representative of the socio-economic diversity in northern Nevada – indicating that free genetic testing is of interest to a broad swath of the population and a powerful motivator for comprehensive population health study research.

## Introduction

Health is the result of the complex and not fully elucidated interaction between genetic, sociodemographic and environmental factors often influenced by the availability or quality of health care [1]. Optimally, these factors need to be integrated in order to develop and apply health care interventions to improve the health of a population.

Depending on the data source used, Nevada ranks in the bottom half of overall health rankings in the United States. The 2016 annual report of America’s Health Ranking for example, ranks Nevada #35 [2]. At the same time, healthcare spending per capita in Nevada is $5,735, 16% below the national average, ranking the state #45 [3]. The majority of residents of Northern Nevada live in Washoe County, which is confounded with high age-adjusted death rates for heart disease, cancer and chronic lower respiratory disease. These death rates exceed those in Nevada and the United States more generally (cumulative deaths in the 3 categories= 501.5 in Washoe County vs. 416.5 in Nevada vs. 385 in the United States per 100,000) [4].

The Renown Institute for Health Innovation (IHI), a newly established public-private partnership between Renown Health (RH) and the Desert Research Institute’s (DRI) Applied Innovation Center (AIC), aims to better understand those factors that contribute to poorer health outcomes in Nevada. This will be achieved by identifying and prioritizing investigations that will lead to insights allowing Nevadans to develop strategies that will improve overall health. The Healthy Nevada Project (HNP) was aimed at two goals. First, to start informing individual Nevadans of their genetic makeup such that they might make better informed decisions to modify their behaviors and improve their health. Since genetic testing as an educational tool is likely biased and restricted to those who can afford the tests (in general selecting for a more affluent population), ours were provided at no cost to participants. Second, to collect comprehensive genetic data on an extensive number of Nevadans that could be used to cross-reference with health, healthcare, social and environmental data to identify specific associations in our local communities and assist in healthcare resource planning. This will be achieved using our expertise in geomapping and understanding the effects of natural and human-induced environmental changes, and the unique computational and storage demands for data hosting, exploration, and visualization. The latter will be used to analyze aggregated electronic health record (EHR) data provided by RH.

## Material and Methods

### Patient Population

According to the United States Census data for 2016, the Washoe County catchment area, at the center of this phase of the study, includes 6,302 square miles and is home to more than 453,000 people (population per square mile 71.9). Of these 22.2% are under the age of 18 years and were excluded from this, first, phase of the study. The study was open to all other members of the community, including, the approximately 13.8% of the population that lives below the Federal Poverty Level.

### Study Appointment Scheduling

23andMe has in place a well-established registration process where individuals can send their saliva from the privacy of their home. However, for the purpose of this study, which included an additional consent to merge de-identified genotype data with de-identified EHR data, an additional registration was required. It was also important to educate each participant thoroughly on the study, its implications, and potential outcomes for Nevada. For these reasons, it was decided to provide participants the opportunity to meet research staff face to face at a physical location for a study participation experience that was well choreographed. This process included a study “meet up” location, a chaperone to the study location, and a video tour of the process. These methods are detailed below.

Investigators utilized Appointment-Plus, a 3^rd^ party HIPAA-compliant commercial appointment provider service [5]. The software allowed study coordinators to manage subject load balance and modify appointment schedules given regular no shows and cancellations.

To schedule an appointment, prospective study participants were guided to a website where they entered basic participant information, including name, birth date, and email address, and subsequently selected a one hour-long time slot to meet research staff. Typically, five, one-hour time slots were offered during each weekday, two, one-hour time slots each Saturday, and one, one-hour time slot each evening from Monday to Friday. Each time slot could include up to 30 participants.

After scheduling an appointment, research coordinators validated the appointment, after which participants were sent an email through Appointment-Plus confirming their registration along with directions to the collection site. In addition, the confirmation email contained the RH-DRI Community Health Project consent document in a Portable Document Format (PDF).

Appointment-Plus generated a reminder for the appointment, emailed two days prior to the scheduled visit.

### Institutional Review Board (IRB) and Consent

All participants were provided with 3 consents. Two consents were provided by 23andMe in electronic format, one to consent for genetic testing and sharing genotype data with the study, the other for DNA bio-banking. The latter consent is optional and does not affect the 23andMe experience or access to genotype data. The third consent form was provided by IHI and provided permission for the de-identified genetic data to be linked to de-identified EHR data from the same study subjects.

#### 23andMe Consent

At the appointment, participants registered with 23andMe via laptop computers available at the sampling site. During this online registration, participants provided 23andMe electronically with the required consents, similar to any other individual who chooses to have their DNA tested by 23andMe. Participants approved the anonymous utilization of their SNP data and also optionally approved for the saliva to be bio banked. Both 23andMe consent forms have been approved by the Ethical and Review Services IRB. Registration with 23andMe is the point where the unique saliva collection tube number is tied to subject demographic information and an email address for 23andMe account creation and further communication.

#### IHI Consent

A copy of the IHI study consent document approved by the University of Nevada Reno IRB was provided electronically to each participant when they received an electronic confirmation of their registration. This gave each participant the opportunity to study the IHI study consent form and prepare questions when they met with study coordinators. During the one-hour appointment, special effort was made to ensure that participants understood the data sharing authorization granted by signing the IHI consent, which would provide the IHI access to the 23andMe genotype data in de-identified form. At their appointment, participants signed the IHI consent form, providing the investigators a permanent record of consent. Each participant received a signed copy of the IHI consent form. data in de-identified form.

A total of 270 participants were provided the option to participate without a formal appointment – the majority of these people were guests on the day of the project launch and 23 were “ambassadors” pre-identified to communicate about the study to the community. These individuals received a consent form immediately prior to providing a saliva sample. However, ample time was provided for them to read and consider the content of the consent form. All participants signed the consent form with a Collaborative IRB Training Initiative (CITI) certified study coordinator available to answer questions and as witness prior to sample collection.

### Acquisition collection kits

23andMe shipped 10,000 saliva collection kits to Renown Regional Medical Center. These kits conform to accepted FDA saliva collection requirements and contain a saliva sample tube (Orogene DX OGD-500.001) with a unique identifier. Each participant is identified by an 11 character mixed-case string to uniquely track study participants. There is no associated protected patient information associated with this participant identifier that is accessible to researchers. Only the IHI Principal Investigator has the link between the participant’s ID and the research data.

### Sampling Location

The sampling was performed in the Renown Regional Medical Center’s wing designated for Research and Education. Each cohort would show up at a public gathering place in the hospital. Research coordinators then checked participants in, arranged them in groups of 10, and escorted them to the study area. The study area was divided into four separate stations: education, 23andMe registration & consent, saliva collection, and sample deposition followed by a final paperwork and quality assurance check by a CITI trained coordinator.

A video was produced to ensure that participants understood and followed the registration and saliva collection process. When cohort members arrived at the saliva collection site they first watched a seven-minute video with instructions for 23andMe registration, and explanation of the RH-DRI study, and saliva collection. The video ensured consistent information was provided to the participants and provided a scalable method of communication to research participants. Next, participants walked to the 23andMe registration station where their test kits were waiting to be registered. There they opened the kits and registered the unique tube number with 23andMe and completed the standard 23andMe online consents. From there, participants went to the saliva collection site to produce the requisite 2 ml sample. This area was partitioned for privacy. After sample collection, participants took their sealed kit to the sample deposition station where they dropped it off and had their paperwork confirmed by a CITI trained research coordinator. Finally, participants were then escorted back to the gathering area.

### English as a Second Language

Recognizing a large Hispanic community in Washoe County, a separate one-day saliva collection event was conducted in Spanish. This required that the consent forms and signage be converted to Spanish and approved by the IRB. Moreover, Spanish-speaking study coordinators were available to coordinate the sample collection/registration in Spanish. A total of 37 Hispanic research participants were recruited during this one-day event.

### Collection in remote areas in Nevada

Nevada has remote areas that are sparsely populated and underserved medically. The desire to include participants from these remote areas and to test our ability to collect at these sites, made us set up offsite saliva collection events in Winnemucca, Yerington, and Fallon. Each event was staffed with 6 study coordinators, who brought all necessary equipment (laptops, saliva collection kits, signage, pens, tape and other supplies) with them in a van.

### Cost to participants

Through a generous donation of the Renown Health Foundation, we were able to offer the DNA testing for free to all participants. The Governor’s Office of Economic Development through a Knowledge Fund grant to the DRI AIC helped fund data scientists.

### Staffing

Support for study execution depended on numerous staff from RH and from DRI. At RH nine FTEs were utilized for research coordination; at DRI four FTEs were utilized to ensure patients could concurrently register for 23andMe and the IHI study. DRI also spearheaded utilization of the appointment system and creation of the two web-based applications for the registration process.

### EHR data

Starting in June 2016, de-identified EHR data spanning more than 10 consecutive years of >1.4 million patients encompassing >26 million provider encounters were transferred to AIC’s Health Insurance Portability and Accountability Act (HIPAA) secure analysis and storage environment. Since then, EHR data are updated quarterly.

### Matching

In order to link subject EHR data to SNP data the participant 23andMe ID was mapped to a participant’s email address, name and date of birth and then de-identified.

### Cardiac Risk Score Calculation

In order to demonstrate the power of IHI’s data, we tested, to demonstrate proof of principle, our ability to replicate a recent study of genetic risk scoring for coronary artery disease using SNP data [6]. Variant calls from the Human Omniexpress-24 array were provided by 23andMe as VCF files. The data were imputed and phased using the Michigan Imputation Server and Eagle v2.3 [7,8]. The reference panel was HapMap2. Reference SNP cluster IDs (rsIDs) were associated with genomic coordinates using SNPTracker [9]. For cardiac risk score calculation, a total of fifty SNPS were examined based on reported findings [6]. Forty-eight of the 50 rsIDs were in our dataset; 31 were genotyped and 17 were imputed. A cardiac risk score was then assessed based on the odds ratio of each SNP, as previously reported [6]. The natural log of the published odds ratio at each SNP was multiplied by the number of risk alleles for each de-identified participant. A final sum was then calculated across all variants. Data were then visualized using a histogram with a bin width of 0.05.

## Results

### Participation and no-show rate

Restricting recruitment to Northern Nevada participants in this phase of our study was primarily guided by geographic convenience with an estimated local population of about 450,000 people in Washoe County. Although the study and announcement by the Governor was covered in local news and press the appointment process was through the Renown Health website. In this early stage, we wanted to demonstrate our ability to develop an infrastructure that would allow for the successful collection of saliva samples from thousands of people in a short time period. Not only were we able to successfully collect saliva from participants in Washoe county, but we also developed an effective ‘mobile sample collection’ mechanism, to obtain DNA from people living in communities in rural Nevada.

We were surprised at the incredible speed with which participants signed on. The study was announced by Governor Brian Sandoval on September 15, 2016 and within two days the registration of 9,700 volunteers for an appointment was complete. Three hundred volunteer slots were reserved for special events. Moreover, ongoing interest for participation was evidenced by a waiting list of more than 4,000 participants. The no-show rate was just over 11% (1,210 of 10,554 online appointments made) and was mitigated by pulling people from a waitlist.

Formal appointments began on September 28, 2016 and were completed on December 21, 2016. Within that time frame there were 69 days of appointments, 17 days off and 9,344 appointments were completed.

### Planning and success

Considerable effort went into planning the public launch. Critical to garnering high-level support in all three organizations was ensuring the Governor of Nevada was interested in participating. The Governor readily agreed to be the first participant in the study on launch day. Dignitaries, board members and employees of both RH and DRI were invited to attend the launch along with members of the press. Additionally, 23 ambassadors were selected from the community to highlight the study. These emissaries were carefully chosen for their visibility in our communities, ranging from a Native American with a high-level position in their tribe, to a radio disc jockey who talked about the study on his radio show.

### Registration and email address

Using a personal email address was a crucial component to the study. First, 23andMe uses the email address as the only mechanism to communicate results with its consumers. When individuals, often a spouse, attempted to utilize a shared email address, the 23andMe registration process, structured to disallow sharing email addresses, would issue a warning “This email is already in use”. Second, the IHI uses the email address provided to identify participants. Occasionally, participants would have used an email to register with 23andMe and a different one when making the appointment for saliva collection. These differences were reconciled so that one email address was used in association with one individual.

### Results from 23andMe

Within approximately eight weeks after donating a saliva sample, each subject received results from 23andMe’s Personal Genome Service, including more than 60 personalized genetic reports that detail their carrier status, genetic mutations which can be passed onto children, for inherited genetic conditions, as well as their traits and ancestry. Earlier this year, the FDA approved 23andMe to provide participants information on their risks to get certain conditions like Parkinson and Late-onset Alzheimer disease. In doing so, 23andMe is the only company with FDA approval to provide genetic health risk reports without prescription. The new information is being made available to participants through 23andMe. Combining genetic research for which results are not immediately available with ‘immediate gratification’ through the 23andMe Personal Genome Service was appreciated by many participants. In fact, we suspect that many participants were swayed to partake because the availability of immediate genetic results, a hypothesis that we intend to test.

### Epidemiology (description of basic demographic data)

#### Race/ethnicity

The racial and Hispanic origin of the broad study region defined as Northern Nevada is compared Northern Nevada to Washoe County, the rest of Nevada and the USA in Table 1. Northern Nevada (population 713,413) is comprised of 11 counties: Carson City, Churchill, Douglas, Elko, Eureka, Humboldt, Lander, Lyon, Pershing, Storey, and Washoe. The majority of residents live in Washoe County (population 453,000). Almost 80% of Northern Nevadans are White and 20% self identify as Hispanic or Latino. Table 2 shows that the study cohort is less diverse than the reported race and ethnicity distribution in Washoe County.

**Table 1.** 2015 Census estimate racial and ethnicity data for Northern Nevada, Washoe County, Nevada and the U.S.

|  | **United States** | **Nevada** | **Northern Nevada** | **Washoe County** |
| --- | --- | --- | --- | --- |
| **White** | 77.10% | 75.70% | 86.70% | 85.10% |
| **Black** | 13.30% | 9.30% | 2.20% | 2.60% |
| **Native American** | 1.20% | 1.60% | 2.90% | 2.20% |
| **Asian** | 5.60% | 8.50% | 4.40% | 6.00% |
| **Pacific Islander** | 0.20% | 0.80% | 0.50% | 0.70% |
| **Other** | 2.60% | 4.10% | 3.30% | 3.50% |
| **Hispanic or Latino** | 17.60% | 28.10% | 22.10% | 23.90% |

U.S. Census Bureau, Population Estimates Program (PEP), Vintage 2015, Accessed 16 May 2017 <https://www.census.gov/quickfacts>.

#### Residence

The residence of the study participants is reported in Supplemental Table 1. In Northern Nevada, RH is the only tertiary care facility between Sacramento, California and Salt Lake City, Utah, attracting patient referrals from surrounding counties. In addition, the Sierra Mountains and Lake Tahoe are vacation destinations that attract a significant number of visitors. This explains the discrepancies noted in Supplemental Table 1: roughly one-third of the RH patient population is from outside of Northern Nevada, while over 98% of study participants are from Northern Nevada. Moreover, for this initial part of the study, the majority of consents were performed at the Renown Regional Medical Center in Reno, NV, self-selecting people living close to the hospital. However, the three rural recruitment events in Fallon, Winnemucca, and Yerington, were highly successful and demonstrated our ability to use a ‘mobile sample collection’ system to obtain saliva for this study from people living in rural Nevada. A total of 501 participants were recruited from these off-site locations (Fallon 160, Winnemucca 168, and Yerington 173). In addition, 551 people from rural Northern Nevada travelled to our central collection center in Reno, resulting in 1,052 participants from outside Washoe County.

#### Age

Age distribution for USA, Nevada, Northern Nevada, and Washoe County are provided in Supplemental Table 2[10]. Of interest the lower % of people under 30 years of age in Northern Nevada as a whole compared to Nevada and the United States (39% versus 41 % and 42% respectively). Washoe County has 42% of people in this age group.

When comparing the age distribution between the Healthy Nevada Project (HNP) participants and the RH EHR patient population we noted several differences (Supplemental Table 3). The HNP participants age distribution is skewed to older participants because individuals 18 years and younger were specifically excluded for this first phase of the study. Second, HNP participants between the ages of 31 and 70 years were over-represented in this cohort, while those over the age of 70 were underrepresented.

**Table 2.** Self-reported ethnicity data for study participants and all Renown patients.

|  | **Study Participants** |  | **All Renown Participants** |  |
| --- | --- | --- | --- | --- |
|  | **Count** | **% of Cohort** | **Count** | **% of Cohort** |
| **White** | 6515 | 88.86% | 986645 | 73.98% |
| **Black** | 91 | 1.24% | 39190 | 2.94% |
| **Native American** | 43 | 0.59% | 15219 | 1.14% |
| **Asian** | 155 | 2.11% | 30786 | 2.31% |
| **Pacific Islander** | 15 | 0.20% | 2969 | 0.22% |
| **Other** | 143 | 1.95% | 44311 | 3.32% |
| **Missing data** | 134 | 1.83% | 78515 | 5.89% |
| **Hispanic or Latino** | 236 | 3.22% | 136086 | 10.20% |

**Table 3.** Location of residence of the study participants.

|  | **Study Participants** |  | **All Renown Patients** |  |
| --- | --- | --- | --- | --- |
|  | **Count** | **% of Cohort** | **Count** | **% of Cohort** |
| **Reno/Sparks** | 6418 | 83.18% | 703656 | 50.52% |
| **All other Washoe County** | 104 | 1.35% | 19319 | 1.39% |
| **All other Northern Nevada** | 1052 | 13.63% | 229877 | 16.50% |
| **All other** | 142 | 1.84% | 440005 | 31.59% |

#### Income/poverty level

The Reno/Sparks urban area has 5 zip codes with greater socio-economic disparity than the rest of Washoe County [4]. Approximately 30% of the population live in these 5 zip codes but represent 42% of the hospital inpatient visits and 54% of the emergency room visits. Supplemental Table 4 demonstrates that the inclusion criteria and our ability to offer the test for free, allowed us to include people from various economic backgrounds. The percentage of participants considered to live below the poverty level did not select only for participants who could afford personal genetic testing. The participant population comprised of approximately 30% who live in the most impoverished Zip Codes of Washoe County indicating no cost participation does provide economic diversity to the cohort.

**Table 4.** 2015 American Community Survey age for Northern Nevada, Washoe County, Nevada and the U.S.

| **Age** | **United States** | **Nevada** | **Northern Nevada** | **Washoe County** |
| --- | --- | --- | --- | --- |
| **Under 5 years** | 6.30% | 6.40% | 5.90% | 6.20% |
| **5 to 9 years** | 6.50% | 6.80% | 6.50% | 6.30% |
| **10 to 14 years** | 6.50% | 6.50% | 6.20% | 6.40% |
| **15 to 19 years** | 6.70% | 6.30% | 6.00% | 6.30% |
| **20 to 24 years** | 7.10% | 6.70% | 6.60% | 7.40% |
| **25 to 34 years** | 13.50% | 14.30% | 13.20% | 14.10% |
| **35 to 44 years** | 12.80% | 13.70% | 12.00% | 12.50% |
| **45 to 54 years** | 13.90% | 13.60% | 13.70% | 13.70% |
| **55 to 59 years** | 6.60% | 6.30% | 7.10% | 6.80% |
| **60 to 64 years** | 5.80% | 5.80% | 6.80% | 6.40% |
| **65 to 74 years** | 7.90% | 8.40% | 10.30% | 8.70% |
| **75 to 84 years** | 4.30% | 3.90% | 4.30% | 3.70% |
| **85 years and over** | 1.90% | 1.30% | 1.50% | 1.50% |

#### Gender

The EHR data population was well balanced in terms of gender reflecting the gender balance in the overall population. The HNP population consisted of many more females (65%) than males (35%)(Supplemental Table 5).

**Table 5.** Age data for study participants and all Renown Patients.

|  | **Study Participants** |  | **All Renown Patients** |  |
| --- | --- | --- | --- | --- |
|  | **Count** | **% of Cohort** | **Count** | **% of Cohort** |
| <18 years | 0 | 0.00% | 143451 | 10.30% |
| 18-30 years | 1052 | 13.63% | 206333 | 14.81% |
| 31-40 years | 1546 | 20.04% | 163433 | 11.73% |
| 41-50 years | 1308 | 16.95% | 172804 | 12.41% |
| 51-60 years | 1476 | 19.13% | 181403 | 13.02% |
| 61-70 years | 1533 | 19.87% | 205401 | 14.75% |
| 71-80 years | 698 | 9.05% | 190203 | 13.66% |
| >80 years | 103 | 1.33% | 129829 | 9.32% |
| Missing Data | 0 | 0.00% | 0 | 0.00% |

**Table 6.** American community survey (2010-4) study of income / poverty level for Northern Nevada, Washoe County, Nevada and the U.S.

|  | United States | Nevada | Northern NV | Washoe County |
| --- | --- | --- | --- | --- |
| **Median Household Income** | \$53,889 | \$51,847 | \$54,150 | \$52,870 |
| **Households Below Poverty Level** | 5.34% | 4.97% | 5.05% | 5.38% |
| **Individuals Age 25+ Below Poverty Level** | 12.01% | 12.17% | 11.27% | 11.89% |

**Table 7.** Sex of study participants and all Renown patients.

|  | Study Participants |  | All Renown Patients |  |
| --- | --- | --- | --- | --- |
|  | Count | % of Cohort | Count | % of Cohort |
| **Female** | 5140 | 66.59% | 685155 | 49.18% |
| **Male** | 2571 | 33.31% | 706272 | 50.70% |
| **Unknown** | 3 | 0.04% | 250 | 0.02% |
| **Missing data** | 5 | 0.06% | 1420 | 0.10% |

### Sample issues

On occasion, there were sample issues – not enough DNA from saliva or other methodological issues – that caused a sample exception in the 23andMe processing pipeline. These were communicated to participants and a new kit was sent to their address of choice. We encountered 21 sample exceptions in our study. The largest problem encountered with the delivery of results to participants was forgotten passwords or other issues accessing established 23andMe accounts. Participants would call the study coordinator number and be directed to 23andMe customer service for privacy purposes (emails and passwords were involved) while an IHI research coordinator followed up to ensure the participant was satisfied with the resolution. Five people dropped out of the study because of dissatisfaction with the experience or concerns with privacy.

### Electronic Healthcare Records (EHR)

Participants consented to have their EHR history at RH, if available, merged with their genetic data in order to study outcomes – this matching was done and then de-identified. Since this genetic study was not restricted to just RH patients, 8180 participants were EHR matched, while the remaining 1520 participants had no EHR data or were unable to be matched.

### Cardiac Risk Scores

Cardiac risk scores were quantified from the SNP data provided by the direct to consumer genetic testing company, 23andMe [6]. The mean risk score of our cohort (3.57) was consistent with other studies (Figure 2). The lowest quartile was 3.208 and upper quartile was 3.918.

**Figure 1.**
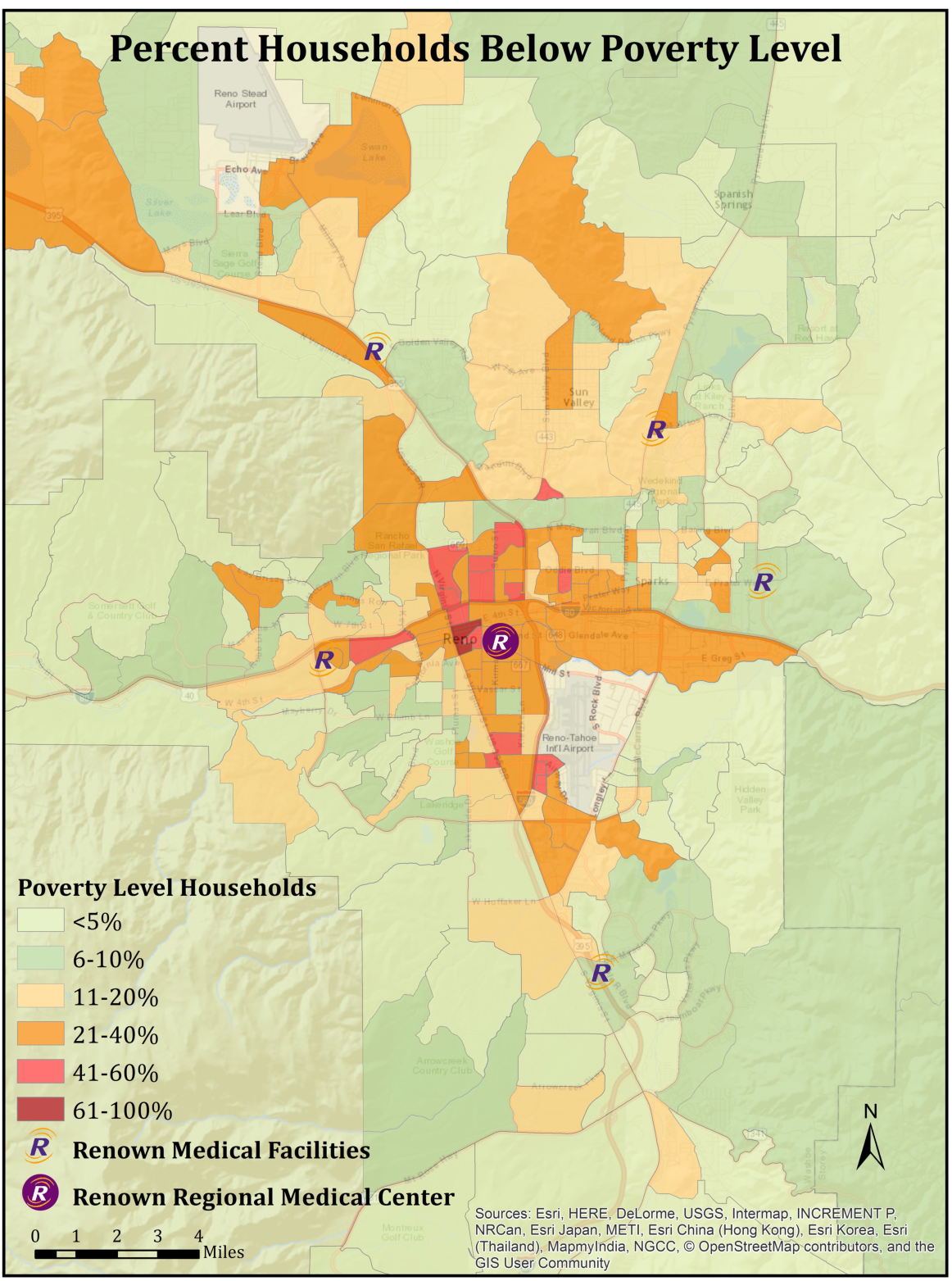
Demographic map of the Reno/Sparks metropolitan area indicating per cent of households in zip code blocks below the poverty level.

**Figure 2.**
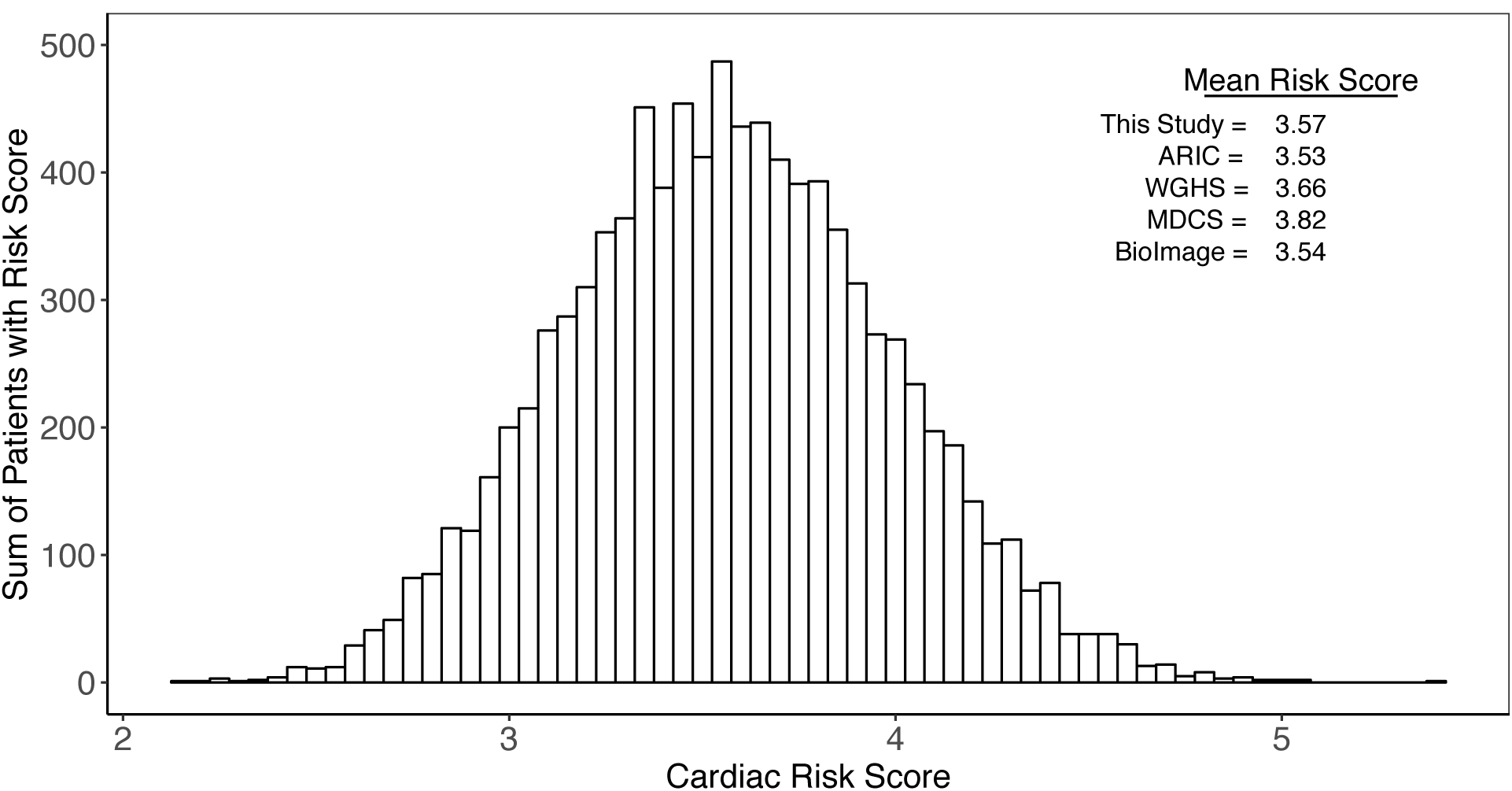
Distribution of cardiac risk scores among the participants in the study. The scores are calculated as in Khera et al. [6] and is described in the text. Risk score means from other studies are indicated [17–20].

## Discussion

The ultimate goal of the Healthy Nevada Project is to recruit 100,000 study participants, representative of Nevada’s population to improve the health of the population through identification of susceptibility for disease, early detection, precision treatment approaches and care delivery research, among many possibilities. This will allow us to link detailed health and healthcare information with comprehensive environmental data and extensive genetic information. Here we report on the first phase of DNA sample recruitment, the acquisition of DNA samples from 9,700 individuals in Northern Nevada, and the proof of principle that our approach can generate the desired data to influence health in our communities.

The enrollment was completed in two days and the consenting and saliva collection was completed in 69 working days. We suggest that the remarkable success can be attributed to a carefully organized marketing campaign and an extremely successful public launch. Prior to the launch, with the input of a local advertising agency, the study was widely advertised ranging from creating a web presence to billboards in the community. Throughout the campaign special attention was given to ensuring consistent messaging. The following factors also contributed to the success of the study. First, garnering sufficient organizational support to conduct the study. This applies in particular to RH that had to make space and personnel available for scheduling sample collections, hosting, informing, and consenting participants, and for collecting and processing the samples. Second, the generous financial support of the Renown Health Foundation was of critical importance since it removed financial barriers to participation. Third, ensuring awareness that the study was taking place in the community and was intended to address a community problem. This involved RH, the DRI, the Governor’s office, and 23andMe. As part of driving home the marketing message, 270 launch attendees were given 23andMe saliva collection kits to begin the study after completing the consent process onsite -- these included 23 ambassadors who were instrumental in helping publicize and share through their networks the potential benefits of participating. Finally, utilizing 23andMe’s well-established processes Foundation was of critical importance since it removed financial barriers to participation. and infrastructure to facilitate sample and data collection, as well as subsequent DNA extraction and processing was a key element of the study. Partnering with 23andMe, who has commoditized the process, allowed for quick, efficient, and quality sample collection and processing.

### Privacy

While potential privacy concerns related to access of the combined (health, healthcare, genetic, social, and environmental) data were clearly dealt with in the consent forms provided, a small number of participants remained concerned with privacy issues associated with the study. Specifically, some participants asked research coordinators if they were required to use their real names, email addresses and birthdates. Incorrect email addresses make account creation with 23andMe impossible while incorrect names and/or birthdates would make matching participants to EHR data impossible. We also noted that less than 10 potential participants, who indicated to have intensively examined the privacy statement on the 23andMe website, opted out at the time of saliva collection. Finally, we have a few participants specifically inquiring whether IHI would share their DNA data with insurance companies. The IHI IRB document expressly states that only DRI will have access to both de-identified DNA and EHR. Consequently, no identifiable genetic data will ever be available to RH, its employees, or any other organization, including insurance companies. This firewall between the hospital system and the research was considered essential to the study. The only way to link the SNP data to EHR record data would be after obtaining specific consent from participants for future studies that would have to be reviewed and approved by an IRB.

### Biases in our convenience sample

There are obvious biases in convenience sampling [11]. The RH population is noticeably on average older; older people use hospital care more frequently than younger (explaining the higher percentages in the EHR data group), while younger people are generally more mobile and able to travel to the study site (explaining the higher percentages in the HNP population). When expanding the IHI study to 100,000 participants special attention will need to be made to recruit more aged individuals to have an accurate reflection of the population.

There are noted sex based biases in volunteer behavioral research [12]. As well, a large majority of the appointments were during the day and it could be that the older population had higher male:female workforce participation. Since during this first phase of the study there had been no attempt to balance the male:female ratio, we have identified a need to gain that balance going forward, when the study expands and will purposefully focus on a large cohort representative of the general Nevada population [13,14].

Though NV is not as ethnically diverse as other states our study population was skewed toward whites and under-represented the Hispanic population. Better bi-lingual advertising and direct communication with the Latino community will be necessary for the future study expansion.

### Using consumer genetics in a population health study

Genotype data collected from consumer platforms coupled with self-reported phenotype information have great potential 15]. However, how these data are communicated and implemented in a healthcare setting is still a challenge. We are in the early stages of developing responsible methods to integrate these metrics into more traditional risk factors such as the American Heart Association healthy lifestyle factors of no current smoking, BMI<30, at minimum weekly physical activity and healthy diet [16]. Yet, it is remarkable that for minimum cost (in healthcare perspectives) we can genotype at population scale and gain important insights into information have great potential weekly physical activity and healthy diet our community genetic health risk.

### Lessons learned

Part of the process was making a video available to each participant that walked him or her through the process and expectations (https://www.youtube.com/watch?v=rWzla8NNhlw). Given the feedback we received, we should have included subtitles for those hard of hearing. In addition, we have decided to develop a video with Spanish subtitles, for obvious reasons. We also believe that the consent form can be further improved. Inability to access results or confusion about the creation of a 23andMe account should be covered in the consent. The cardboard boxes that the kits arrive in are superfluous waste when utilizing almost 10,000 in less than 3 months. They should be eliminated. Future sampling events will have to address the incongruity between the diversity of the extant population and those who choose to participate if we are to serve the health needs of all members of our communities.

## Conclusion

A preliminary pilot population health study was initiated and the first phase was to collect saliva and genotype from approximately 10,000 northern Nevadans. This was completed in a catchment area surrounding Reno, NV where a single tertiary care facility treats ∼70% of the patient population. The participant population was skewed to female and was less racially diverse than the population. The genetic test was administered free-of-charge and the patient population was representative of the socio-economic diversity in northern Nevada. More specific results for geographic, environmental, social and genetic patterns will be elucidated as the study proceeds. The intent here was to highlight both the process and aggregate results of the participants as compared to the population from which that sample was obtained.

## Declarations

### Ethics approval

This study was approved by the University of Nevada, Reno Institutional Review Board (956068-8). Each participant provided written, informed consent.

### Consent for publication

Not applicable.

### Availability of data and materials

De-identified participant demographic data and SNP frequencies used for the cardiac risk score calculations are available from the corresponding author on reasonable request.

### Authors’ contributions

JG, CG and TS conceived of the study. MH, JG, JM, BL made substantial contributions to design. JM, BL, HR, SR, RR made substantial contributions to data analysis. JG, JM, CR, CG, MC, TS made substantial contributions to interpretation of data. JG, JM, CR, CG, MC, TS made substantial contributions drafting the manuscript. All authors made substantial contributions analyzing data and revising the manuscript.

## Acknowledgements

The authors wish to thank all of the people in the Renown Office of Research and Education who contributed to making this study a success. We thank the countless volunteers and guides. We thank the Renown IT staff and the DRI IT staff for support. We thank 23andMe and especially the 23andMe GSR team for help during the study recruitment.

---

**Supplemental Table 1.** Location of residence of the study participants.

|  | Study Participants |  | All Renown Patients |  |
| --- | --- | --- | --- | --- |
|  | Count | % of Cohort | Count | % of Cohort |
| **Reno/Sparks** | 6418 | 83.18% | 703656 | 50.52% |
| **All other Washoe County** | 104 | 1.35% | 19319 | 1.39% |
| **All other Northern Nevada** | 1052 | 13.63% | 229877 | 16.50% |
| **All other** | 142 | 1.84% | 440005 | 31.59% |

**Supplemental Table 2.** 2015 American Community Survey age for Northern Nevada, Washoe County, Nevada and the U.S.

| Age | United States | Nevada | Northern Nevada | Washoe County |
| --- | --- | --- | --- | --- |
| **Under 5 years** | 6.30% | 6.40% | 5.90% | 6.20% |
| **5 to 9 years** | 6.50% | 6.80% | 6.50% | 6.30% |
| **10 to 14 years** | 6.50% | 6.50% | 6.20% | 6.40% |
| **15 to 19 years** | 6.70% | 6.30% | 6.00% | 6.30% |
| **20 to 24 years** | 7.10% | 6.70% | 6.60% | 7.40% |
| **25 to 34 years** | 13.50% | 14.30% | 13.20% | 14.10% |
| **35 to 44 years** | 12.80% | 13.70% | 12.00% | 12.50% |
| **45 to 54 years** | 13.90% | 13.60% | 13.70% | 13.70% |
| **55 to 59 years** | 6.60% | 6.30% | 7.10% | 6.80% |
| **60 to 64 years** | 5.80% | 5.80% | 6.80% | 6.40% |
| **65 to 74 years** | 7.90% | 8.40% | 10.30% | 8.70% |
| **75 to 84 years** | 4.30% | 3.90% | 4.30% | 3.70% |
| **85 years and over** | 1.90% | 1.30% | 1.50% | 1.50% |

**Supplemental Table 3.**
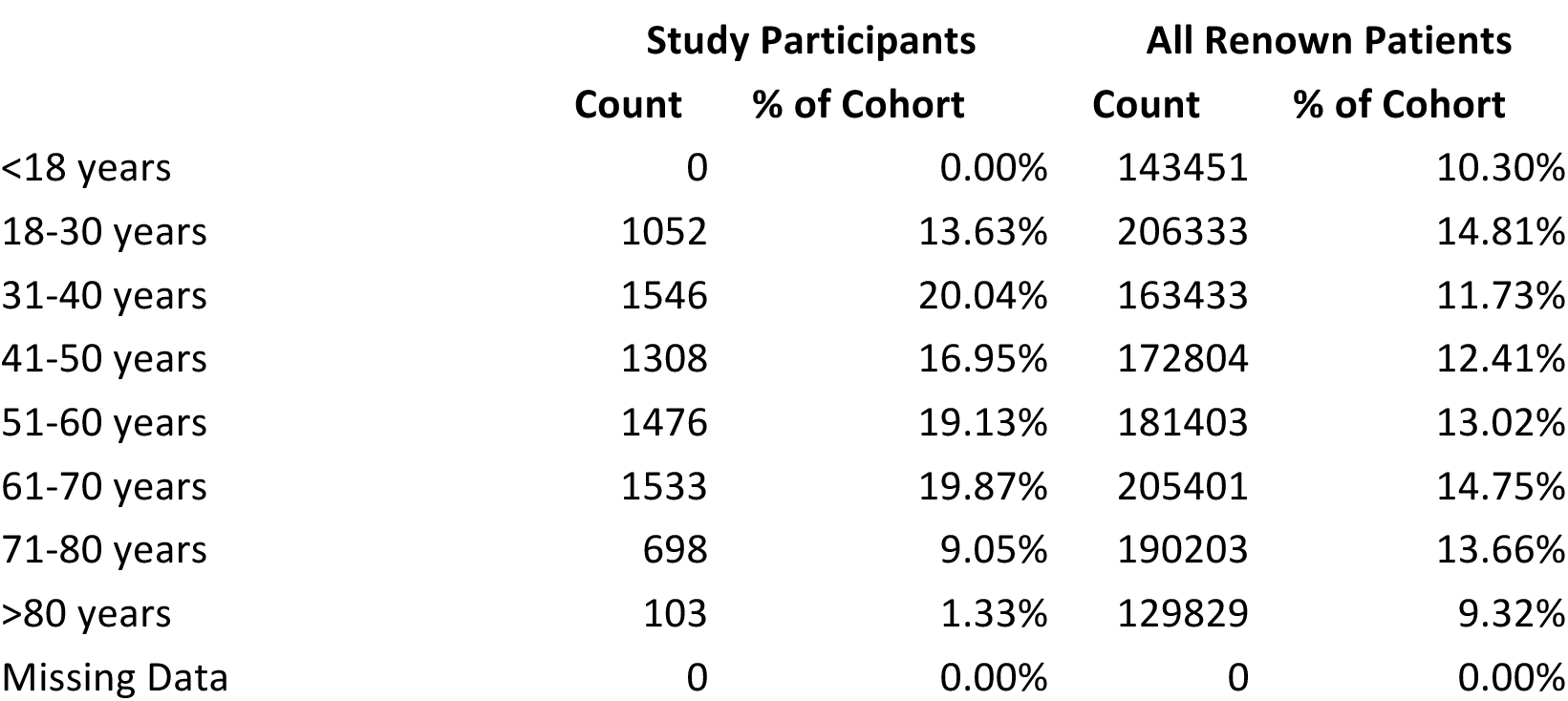
Age data for study participants and all Renown Patients

**Supplemental Table 4.** American community survey (2010-4) study of income / poverty level for Northern Nevada, Washoe County, Nevada and the U.S.

|  | **United States** | **Nevada** | **Northern NV** | **Washoe County** |
| --- | --- | --- | --- | --- |
| **Median Household Income** | \$53,889 | \$51,847 | \$54,150 | \$52,870 |
| **Households Below Poverty Level** | 5.34% | 4.97% | 5.05% | 5.38% |
| **Individuals Age 25+ Below Poverty Level** | 12.01% | 12.17% | 11.27% | 11.89% |

**Supplemental Table 5.** Sex of study participants and all Renown patients.

|  | **Study Participants** |  | **All Renown Patients** |  |
| --- | --- | --- | --- | --- |
|  | **Count** | **% of Cohort** | **Count** | **% of Cohort** |
| **Female** | 5140 | 66.59% | 685155 | 49.18% |
| **Male** | 2571 | 33.31% | 706272 | 50.70% |
| **Unknown** | 3 | 0.04% | 250 | 0.02% |
| **Missing data** | 5 | 0.06% | 1420 | 0.10% |

